# The HSP-90 co-chaperone CDC-37 regulates oomycete immunity in *C. elegans* through a membrane-associated pseudokinase

**DOI:** 10.64898/2026.07.31.741986

**Authors:** Franziska Trusch, Michalis Barkoulas

## Abstract

Innate immune responses depend on rapid and tightly regulated signalling to ensure effective defence against invading pathogens. In *Caenorhabditis elegans*, the oomycete recognition response (ORR) is a transcriptional programme that provides protection against lethal infections by oomycetes, such as *Myzocytiopsis humicola*. The protein tyrosine kinase OLD-1 and the pseudokinase FLOR-1 are essential for activation of this immune response, however, the underlying molecular mechanism is poorly defined. Here, we show that FLOR-1 is regulated by the kinase-specific HSP- 90 co-chaperone CDC-37. Following tandem mass tag-based proteomics, we reveal that FLOR-1 interacts with CDC-37 and that disruption of the CDC-37/HSP-90 complex abolishes ORR gene induction. CDC-37 is required for membrane association of FLOR-1 and mediates its pathogen recognition-induced relocalisation from the plasma membrane to the cytosol, where downstream immune signalling is initiated. Surprisingly, the related active protein tyrosine kinase OLD-1 functions independently of CDC-37. Functional dissection of FLOR-1 localisation demonstrates that membrane-associated FLOR-1 is required for pathogen recognition, whereas cytosolic FLOR-1 is sufficient to activate ORR gene expression. Together, these findings uncover an unexpected role for the CDC-37/HSP-90 complex in regulating innate immunity likely through spatial control of a membrane-associated pseudokinase and reveal a chaperone-mediated mechanism that links pathogen detection to immune activation.

## INTRODUCTION

All organisms constantly face biotic stress, and immune responses are essential for overcoming exposure to pathogens. The nematode *Ceanorhabditis elegans* is a powerful model organism for investigating immune responses to eukaryotic, prokaryotic as well as viral pathogens owing to its ease of genetic manipulation and experimentation, its transparent body suitable for imaging, and the significant overlap with biological pathways in humans (*1, 2*). We previously discovered oomycetes as natural pathogens of *C. elegans* (*3*). Most oomycetes are pathogenic and cause tremendous losses for the agri- as well as aquaculture industries globally by infecting important food sources such as potatoes, soybeans and fish (*4*). So, *C. elegans* is an easily accessible laboratory organism to study host-oomycete interactions.

In contrast to humans, *C. elegans* relies exclusively on non-specialised immune cells such as epithelial cells of the epidermis to trigger an immunity response. Among the canonical immune signalling pathways in nematodes are the p38/MAPK cascade (p38 Mitogen-Activated Protein Kinase) or the MAPK/ERK pathway upregulated upon infection with bacterial and fungal pathogens (*2, 5*). Furthermore, the DAF-2/insulin-like pathway commonly known to regulate ageing, also facilitates survival of laboratory-induced infections with clinically relevant human pathogens such as *Pseudomonas aeroguinosa* (*6*) and *Cryptococcus neoformans* (*7*).

In the case of oomycete infections, the damage-independent oomycete recognition response (ORR) is triggered upon contact with the pathogen or exposure to a non-infectious pathogen extract (*8*). As part of this transcriptional programme, a set of *chitinase-like* (*chil*) genes, amongst others, are induced in the epidermis resulting in modification of the cuticle and enhanced resistance to oomycete infection with *Myzocytiopsis humicola* (*3*). In a recent forward genetic screen aimed at dissecting the machinery underlying pathogen recognition, the membrane- associated FLOR-1 and OLD-1 have been identified as crucial for activating the ORR (*9*). Both proteins belong to the KIN-16 family of receptor-like RTKs, a nematode-specific expansion of the kinome that includes members whose functions and regulation are poorly understood (*10, 11*). Interestingly, while OLD-1 is predicted to be an active kinase, FLOR-1 is a pseudokinase lacking many key motifs that are essential for kinase activity (*9*). Most kinases of the canonical immune signalling pathways in nematodes are cytosolic serine/threonine or dual specificity kinases, underscoring the need to expand our understanding of tyrosine protein kinases in immunity.

About 60% of the human kinome involved in a broad variety of cellular processes are regulated by the CDC-37/HSP-90 complex (*12, 13*). CDC-37 (cell division cycle 37) is a highly conserved, essential integral component of the chaperone network, specifically recruiting unstable client kinases to HSP-90 for protection from degradation but also to reduce unwanted basal activity (*14–16*). CDC37 directly interacts with kinases via its N-terminus in the absence of Hsp-90 and subsequently targets Hsp90 through a central binding domain, forming a tertiary complex that propagates proper folding, stabilisation and activation of the kinase (*17–20*). Later, the client kinase is released which usually requires the Ser/Thr protein phosphatase PPH-5 (mammalian PPP5C), which removes regulatory phosphorylation and restores the kinase as well as the CDC-37/HSP-90 complex to its basal conformation prepared to accept other kinases (*21*). Subsequently, the mature kinase contributes to downstream signalling or when proper folding is lost is then recruited by the HSP-70 chaperone system and prepared for degradation following ubiquitination (*14*). While the role of CDC-37 in facilitating the activity of cytosolic protein kinases is well reported (*12, 14, 22, 23*), the contribution of CDC-37 to stabilising membrane-associated kinases or even pseudokinases is less understood.

Here, we sought to extend our understanding of how epidermal kinases regulate the ORR. We report on the protein interactome of FLOR-1 identified by Tandem Mass Tag (TMT) proteomics revealing an interaction with CDC-37. We demonstrate that the membrane association of FLOR-1 and, concomitantly, the induction of the immune response upon pathogen extract treatment are dependent on the CDC-37/HSP-90 complex. Furthermore, upon exposure to pathogen extract, FLOR-1 is relocalised from the membrane to the cytosol where it induces components of the ORR. While the role of the CDC-37/HSP-90 complex is to facilitate the stability and activity of protein kinases, in this case the effect of CDC-37 appears to be specific to the pseudokinase FLOR-1 as opposed to the active protein tyrosine kinase OLD-1. This report provides a paradigm of how CDC-37/HSP-90 can regulate immune responses through a membrane-associated pseudokinase.

## RESULTS

### Investigating the FLOR-1 interactome after exposure to M. humicola extract

To elucidate the regulation of the oomycete recognition response (ORR) mediated by the pseudokinase FLOR-1, tandem mass tag (TMT) proteomics was employed to identify FLOR-1 interacting proteins (Fig. 1A). In order, to reproducibly trigger the ORR and bypass the spatial and temporal variability associated with natural infections, animals were exposed to an extract of *M. humicola* previously shown to be sufficient to induce the ORR (*8*) and collected in large, synchronised populations given the low protein levels of FLOR-1::GFP (*9*). FLOR-1::GFP was immunoprecipitated from extract treated and non-treated animals. No consistent reduction in band intensity was observed indicating extract treatment does not result in FLOR-1::GFP degradation. In addition, the band pattern and height are comparable between treated and non-treated conditions and therefore, it is unlikely that extract treatment causes posttranslational modifications of the pseudokinase (Fig. 1B).

**Figure 1.**
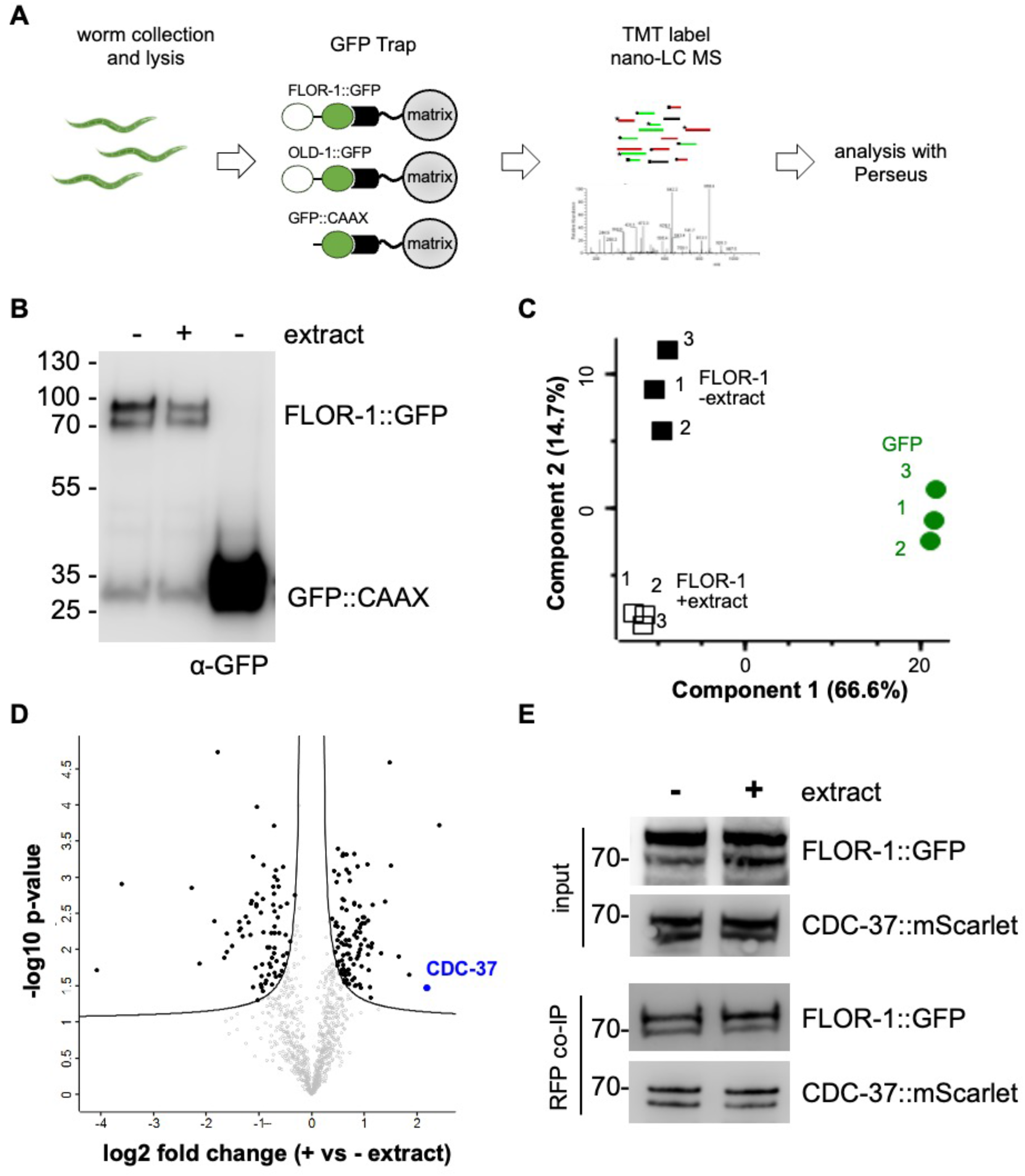
TMT-based proteomic analysis reveals FLOR-1 interactors upon *M. humicola* extract exposure. **(A)** Overview of the TMT proteomics workflow used to identify proteins interacting with FLOR-1::GFP following exposure to *M. humicola* extract. After immunoprecipitation and TMT labelling, samples were analysed by LC– MS/MS to quantify changes in the FLOR-1 interactome. **(B)** Representative immunoblot showing co-immunoprecipitated FLOR-1::GFP with (+) and without extract (-) exposure, and the GFP::CAAX control used for TMT proteomics. Molecular weights are indicated in kilodaltons (kDa). **(C)** Principal component analysis showing clear separation of FLOR-1::GFP interactomes with (empty squares) and without extract (black squares) in comparison to the GFP::CAAX control (green circles), indicating global proteomic changes in the presence of *M. humicola* extract. Each point represents one biological replicate, and the proportion of the total variation for each principal component is shown in parentheses. **(D)** Volcano plot visualising log_2_ fold change and statistical significance of FLOR-1::GFP interactors upon extract treatment (+) compared to no extract (-). A total of 293 proteins showed significant differential enrichment (FDR < 0.05, filled circles) amongst 983 identified proteins in total. The co-chaperone CDC-37 studied here is highlighted in blue. **(E)** Co-immunoprecipitation in animals expressing FLOR-1::GFP and CDC-37::mScarlet using RFP-trapping confirms the interaction identified by TMT proteomics. Extract exposure (+) does not alter the band pattern or affinity between FLOR-1::GFP and CDC-37::mScarlet compared to no extract (-). Molecular weights are indicated in kilodaltons (kDa).

For analysis of TMT proteomics proteins were normalised to membrane bound GFP::CAAX controls which allows the relative quantification of potential interactors. Principal component analysis of the data set confirmed tight clustering of biological triplicates and a clear separation between extract treated and non-treated samples as well as GFP controls (Fig. 1C). Initially, FLOR-1::GFP without extract treatment was compared to GFP::CAAX, resulting in 477 proteins with significant differential enrichment out of 983 protein hits (Fig. S1A, Table S1). Gene set enrichment analysis indicated that proteins binding to FLOR-1::GFP under normal conditions amongst others are involved in signalling, proteolysis, chaperoning and trafficking (Fig. S1B). Subsequently, to narrow down which of those proteins are relevant for the response to *M. humicola* extract, the interactome of FLOR-1::GFP with and without extract was compared. We found that 100 interactions with FLOR- 1::GFP were strengthened upon extract exposure, while 73 were reduced during treatment with extract (Fig. 1D, Table S2). Surprisingly, despite protein levels of the protein tyrosine kinase OLD-1 being dependent on the pseudokinase FLOR-1 (Fig. S1C and S1D), no interaction between FLOR-1 and OLD-1 was detected (Table S1).

Consistently, TMT proteomics of OLD-1::GFP did not confirm a significant interaction with endogenous FLOR-1 (Table S3). In addition, co-immunoprecipitation with animals co-expressing FLOR-1::GFP and OLD-1::mCherry also failed to detect an interaction by either GFP or RFP pull downs, although reciprocal immunoprecipitations confirmed sufficient expression of both proteins (Fig. S1E).

To narrow down key FLOR-1::GFP interactors, a targeted RNAi screen was performed prioritising candidates involved in signal transduction, protein sorting and stability as well as other protein kinases that might potentially form a signalling cascade (Table S4). Immune signalling was assessed based on the extract-mediated induction of two ORR reporters, a transcriptional *chil-27p*::GFP as well as a translational B0507.8::GFP. Among all candidates tested, *cdc-37* emerged as important because *cdc-37* RNAi greatly reduced induction of both reporters upon exposure to *M. humicola* extract (Fig. S1F). In addition, co-immunoprecipitation of CDC-37::mScarlet and FLOR-1::GFP confirmed the interaction (Fig. 1E) indicating that CDC-*37* is required for the ORR likely through its association with the pseudokinase FLOR-1. CDC-37 is a part of a molecular chaperone system involved in the stabilisation and activation of protein kinases. In *C. elegans*, CDC-37 has been previously implicated in the activation of MBK-2, a dual specificity tyrosine-phosphorylation-regulated kinase involved in marking proteins important for the oocyte-to-zygote transition for degradation (*24*). Although, *mbk-2* is significantly expressed beyond the germline in ORR-relevant tissues including the epidermis, intestine as well as neurons (*25, 26*), *mbk-2* RNAi did not impair the induction of the two ORR genes in response to extract (Fig. S1F), indicating that CDC-*37* may be required for the activation of the ORR through its association with the pseudokinase FLOR-1 and not MBK-2.

### The CDC-37/HSP-90 complex acts on the pseudokinase FLOR-1 but not the protein kinase OLD-1

CDC-37 recruits client protein kinases to the molecular chaperone HSP-90, forming a ternary complex which is then resolved by removing activating phosphate groups by PPH-5 (*21*). Hence, the contribution of other CDC-37 complex components to the ORR was investigated by examining the effects of *cdc-37* and *hsp-90* RNAi as well as a *pph-5(ok3498)* deletion mutant on the *chil-27p*::GFP response to *M. humicola* extract. Similar to *cdc-37* RNAi, knockdown of *hsp-90* by RNAi strongly reduced the number of *chil-27p*::GFP positive animals upon extract exposure (Fig. 2A and 2B). In *pph-5*(-) mutants, the *chil-27p*::GFP response to extract was completely abolished (Fig. S2A and S2B). However, the simultaneous loss of the non-extract inducible *col- 12p*::DsRed marker carried in the background suggested potential transgene silencing. In fact, qPCR analysis of another ORR gene B0507.8 confirmed that endogenous transcript levels still increased upon continuous extract exposure in *pph-5*(*-*) mutants, comparable to wild-type (Fig. S2C). In contrast, the effect of *cdc- 37* RNAi was also reflected in the reduced transcript levels of other ORR genes upon extract treatment, such as *Y71H2AR.2* and *B0507.8* (Fig. 2C). We note that our focus was on *cdc-37* RNAi because undiluted *hsp-90* RNAi results in L1 arrest and hence, the effect of diluted RNAi is required, which is more difficult to control.

**Figure 2.**
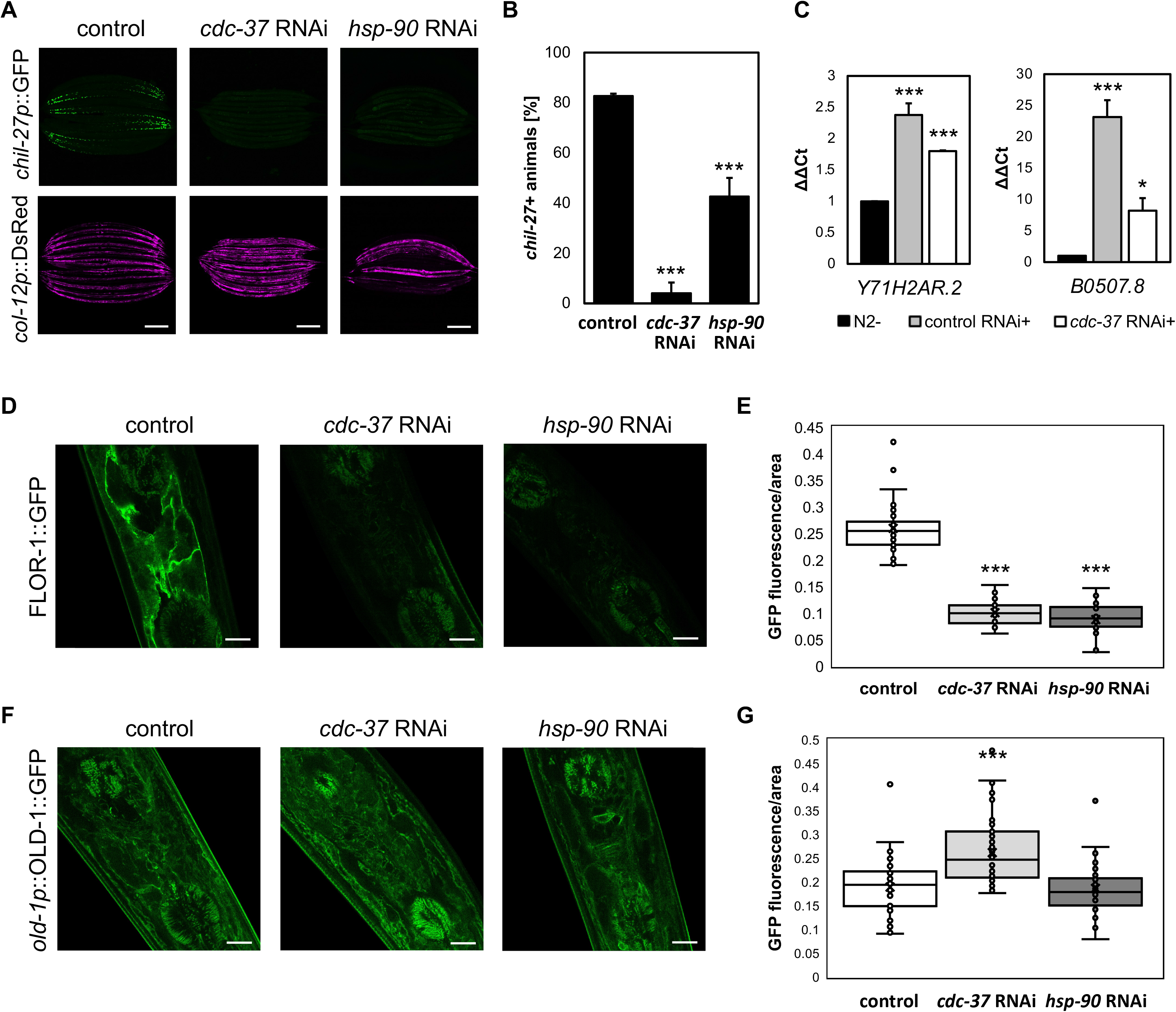
HSP-90 and its co-chaperone CDC-37 are required for the *chil-27p*::GFP response to *M. humicola* extract and regulate FLOR-1 levels. **(A)** Representative images of *chil-27p*::GFP expression in day 1 adults of the following generation subjected to *cdc-37* and *hsp-90* RNAi following 24 h with *M. humicola* extract exposure. The *chil-27p*::GFP response is significantly reduced by both RNAi treatments. **(B)** Quantification of *chil-27p*::GFP response to extract after *cdc-37* and *hsp-90* RNAi as shown in (A), (n=30 per condition, biological triplicates). **(C)** qPCR quantification of additional ORR genes (Y71H2AR.2, B0507.8) upon extract exposure (+) for 72 h of animals on control and *cdc-37* RNAi normalised to untreated N2 (-) suggests that *cdc-37* probably affects multiple genes of the ORR (biological triplicates). **(D)** Representative images showing reduced membrane localization of FLOR-1::GFP in the anterior head region of day 1 adults after *cdc-37* and *hsp-90* RNAi. Insets for *cdc-37* and *hsp-90* RNAi display the same image with enhanced contrast. **(E)** Quantification of FLOR-1::GFP fluorescence per area anterior in the head of day 1 adults as shown in (D), (n=30 per condition). **(F)** In contrast to FLOR-1, representative images show an increase or no effect on the membrane localization of OLD-1::GFP in the anterior head region of day 1 adults after *cdc-37* and *hsp-90* RNAi, respectively. **(G)** Quantification of OLD-1::GFP fluorescence per area anterior in the head of day 1 adults as shown in (F), (n=40 per condition). Scale bars: (A) = 200 μM, (D, F) = 10 μM. Bars in (B) and (C) are mean±SD. (B) = \**p* < 0.05, (B, C, E, G) = \*\*\**p* < 0.001, unpaired t-test.

Since FLOR-1 and CDC-37 were found to be associated, the effect of depletion of single components of the CDC-37/HSP-90 complex on FLOR-1::GFP protein levels was investigated (Fig. 2D and 2E). Indeed, *cdc-37* and *hsp-90* RNAi significantly reduced membrane-associated FLOR-1::GFP levels. In contrast, *pph-5*(*-*) animals showed no effect on FLOR-1::GFP (Fig. S2D and S2E), which is consistent with the notion that the observed effect on induction of the ORR markers is more likely due to transgene silencing rather than direct regulation of FLOR-1 stability or localisation. For comparison we also quantified membrane-associated protein levels of the active tyrosine kinase OLD-1::GFP upon *cdc-37* and *hsp-90* RNAi as well as in *pph-5*(*-*) mutants. While *hsp-90* RNAi had no effect on OLD-1::GFP levels, both *cdc- 37* RNAi and *pph-5*(*-*) mutants showed increased membrane accumulation of OLD- 1::GFP, opposite to the effect observed for FLOR-1 (Fig. 2F, 2G, S2F and S2G). In addition, TMT proteomics of OLD-1::GFP did not detect any significant interaction with CDC-37 (Table S3). Taken all together, these results demonstrate that the CDC-37/HSP-90 complex is crucial for the activation of the ORR specifically acting on the pseudokinase FLOR-1 and independently of OLD-1.

### CDC-37 regulates FLOR-1 relocalisation in response to M. humicola extract treatment

Endogenous FLOR-1 protein abundance is low, which makes it difficult to determine whether the reduction or FLOR-1::GFP fluorescence intensity observed upon *cdc- 37/hsp-90* RNAi reflects protein degradation or relocalisation from the membrane. However, we reasoned that the lack of change of band intensities or pattern of FLOR-1::GFP upon extract exposure renders protein degradation as a cause of the apparent reduction unlikely (Fig. 1B and 1E). Furthermore, published RNAseq experiments have not revealed any significant changes of *flor-1* expression during oomycete infection nor upon treatment with pathogen extract (*8, 9*). To visualise potential subcellular changes of FLOR-1, FLOR-1::GFP was expressed under a strong epidermal promoter (*dpy-7p*). Consistent with the endogenous FLOR-1::GFP signal (Fig. 2D and 2E), *cdc-37* RNAi on overexpressed FLOR-1::GFP also reduced total GFP intensity (Fig. 3A and 3B). Although, an increase of the mean cytosolic fraction of FLOR-1::GFP was observed this effect was not statistically significant (Fig. 3C). Furthermore, prolonged exposure to *M. humicola* extract similarly reduced membrane-associated FLOR-1::GFP protein levels (Fig. 3D and 3E) while simultaneously the ratio of cytosolic to total FLOR-1::GFP significantly increases, suggesting a relocalisation from the membrane to the cytosol in the presence of extract (Fig. 3F).

**Figure 3.**
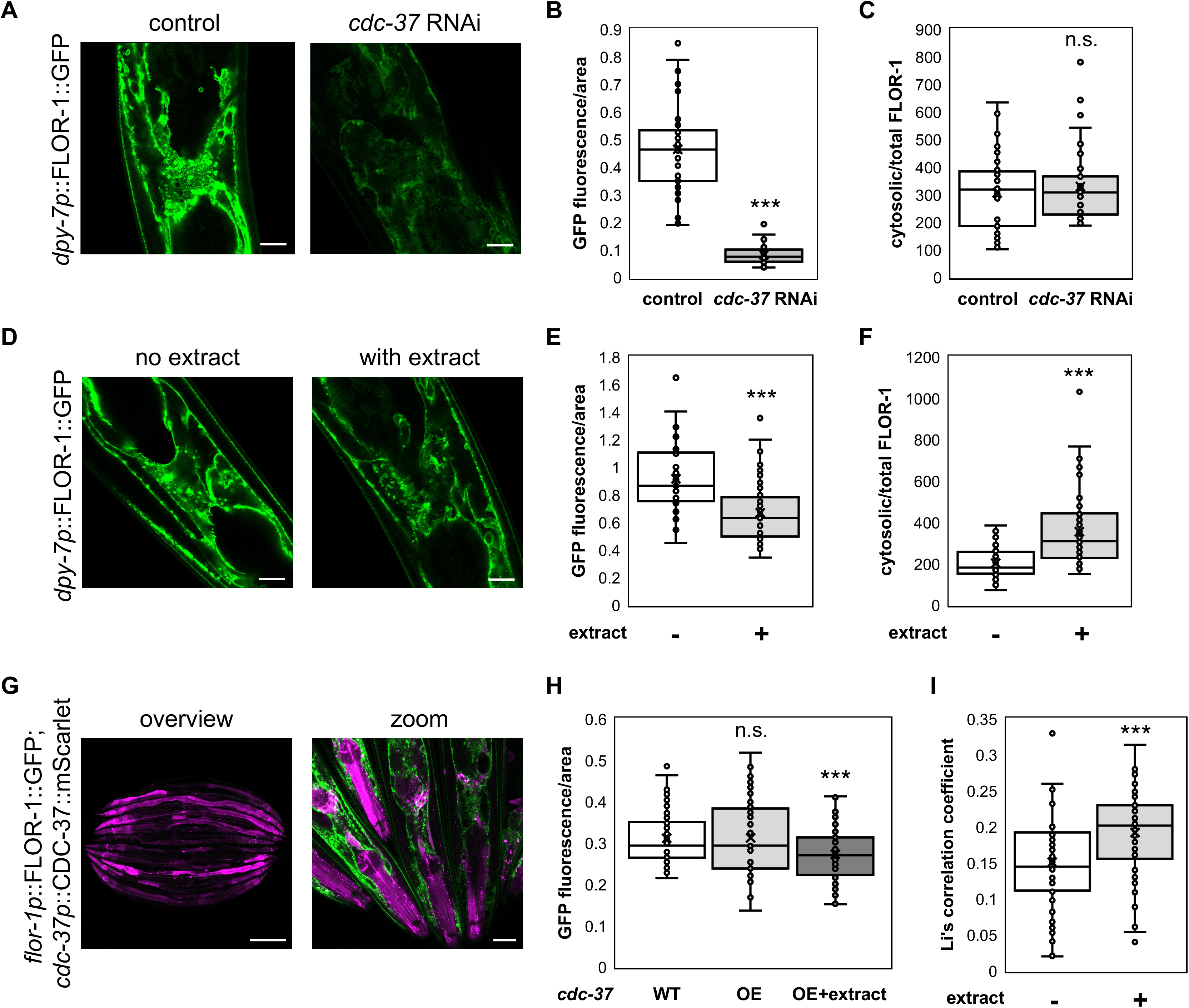
FLOR-1::GFP relocalises from the membrane to the cytosol upon exposure to *M. humicola* extract. **(A)** Representative images showing *dpy-7p*::FLOR-1::GFP localisation anterior in the head of day 1 adults subjected to control and *cdc-37* RNAi. **(B)** Quantification of *dpy-7p*::FLOR-1::GFP intensity per area anterior in the head of day 1 adults as show in (A), (n=40 per condition). **(C)** Ratio of cytosolic *dpy-7p*::FLOR-1::GFP to total *dpy-7p*::FLOR-1::GFP intensity per area (anterior in the head) of day 1 adults as show in (A) upon HT115 or *cdc-37* RNA*i* (n=40 per condition). **(D)** Representative images showing *dpy-7p*::FLOR-1::GFP localisation anterior in the head of day 1 adults with and without exposure to *M. humicola* extract for 72 h supporting relocalisation of FLOR-1 upon extract exposure. **(E)** Quantification of *dpy-7p*::FLOR-1::GFP intensity per area anterior in the head of day 1 adults as show in (D) with (+) and without (-) extract treatment, (n=40 per condition). **(F)** Ratio of cytosolic *dpy-7p*::FLOR-1::GFP to total *dpy-7p*::FLOR-1::GFP intensity per area (anterior in the head) of day 1 adults as show in (D) with (+) and without (-) extract treatment, (n=40 per condition). **(G)** Representative images of membrane-associated FLOR-1::GFP and ubiquitous *cdc-37p*::CDC-37::mScarlet. **(H)** Quantification of FLOR-1::GFP intensity per area anterior in the head of day 1 adults in wild-type and upon CDC-37 overexpression as shown in (G) and upon CDC-37 overexpression as well as continuous extract treatment. Overexpression of CDC-37::mScarlet does not alter membrane FLOR-1::GFP levels, only after extract exposure, (n=40 per condition). **(I)** Graphical presentation of Li’s correlation coefficient between FLOR-1::GFP and CDC-37::mScarlet colocalisation with (+) and without (-) extract treatment. Scale bars: (A, D) = 10 μM, (G) = 200 μM and 20 μM for zoom. (B, C, E, F, H, I) = \*\*\**p* < 0.001, unpaired t-test.

To assess whether increased CDC-37 abundance is sufficient to alter FLOR-1 localisation, FLOR-1::GFP was quantified in animals overexpressing CDC- 37::mScarlet under its own promoter (Fig. 3G and 3H). Interestingly, CDC-37 overexpression neither affected total FLOR-1::GFP levels nor its membrane localisation under basal conditions, but a significant reduction of FLOR-1::GFP was found upon combined CDC-37 overexpression and extract exposure (Fig. 3H). In addition, the correlation of fluorescence intensity between FLOR-1::GFP and CDC- 37::mScarlet increased by 38% upon extract exposure (Li’s correlation coefficient 0.145 vs 0.2 upon extract exposure, n= 50, *p*\*\*\* < 0.001 with Mann-Whitney test, Fig. 3I), which is in line with TMT proteomics showing increased CDC-37/FLOR-1 interaction upon extract treatment (Fig. 1D). Unfortunately, a similar comparison with OLD-1 was not possible due to undetectable OLD-1::GFP levels in a strain co-expressing CDC-37::mScarlet.

To further investigate the effect of a potential activation of CDC-37 on FLOR- 1, a *cdc-37(ax2001)* gain-of-function (GoF) mutant (Leu221Phe; (*27*)) was tested to determine whether enhanced CDC-37 activity is sufficient to influence FLOR-1 relocalisation. Like CDC-37 overexpression, *cdc-37(ax2001)* GoF mutants exhibited normal FLOR-1::GFP distribution in the absence of extract (Fig. S3A and S3B), and similarly, OLD-1::GFP also remained unchanged (Fig. S3C and S3D). However, *cdc- 37(ax2001)* GoF animals showed an increased *chil-27p*::GFP response upon continuous *M. humicola* extract exposure at high dilutions (Fig. S3E and S3F), which is also supported by increased transcript levels of other ORR genes upon extract treatment (Fig. S3G). This suggests that activated CDC-37 can strengthen the response to pathogen extract. Taken together, we hypothesise that CDC-37 facilitates the extraction of pathogen-triggered FLOR-1 from the membrane with a persisting association upon extract exposure.

### Membrane-associated and cytosolic FLOR-1 possess distinct functions

The relocalisation of a transmembrane pseudokinase to the cytosol without subsequent degradation, prompted the dissection of the potentially different functions of membrane-associated and cytosolic FLOR-1. To mimic cytosolic FLOR- 1, the signal peptide as well as transmembrane helix (1-78 amino acids) were removed through genome editing, leaving only the pseudokinase domain (Fig. 4A). As predicted, the truncated FLOR-1(79-466)::GFP displayed diffuse cytosolic localisation, lacking the membrane enrichment of the full-length protein (Fig. 4B). Interestingly, in the absence of pathogen extract, basal activation of *chil-27p*::GFP (Fig. 4C and S4A) as well as upregulation of other ORR genes (Fig. 4D) was observed with the truncated FLOR-1(79-466)::GFP, but not with the full-length FLOR-1::GFP. However, in contrast to full-length FLOR-1::GFP, exposure to *M. humicola* extract did not strengthen the activation of *chil-27p*::GFP in the case of FLOR-1(79-466)::GFP (Fig. 4C). Hence, full extract response requires membrane-associated FLOR-1, whereas cytosolic FLOR-1 can also initiate downstream signalling.

**Figure 4.**
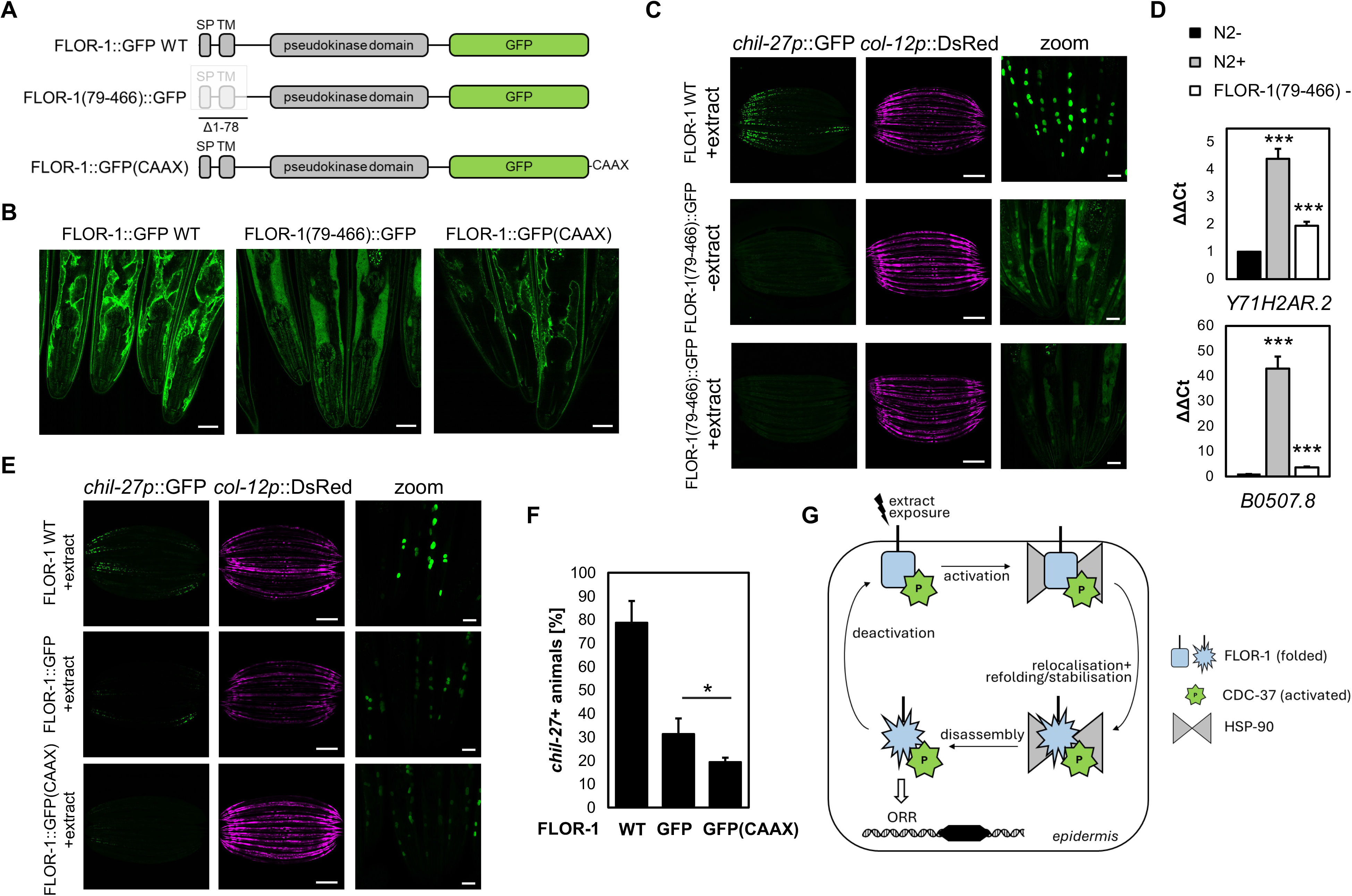
Cytosolic FLOR-1 kinase exhibits basal activity in the absence of *M. humicola* extract yet does not mediate response to extract. **(A)** Schematic diagram of FLOR-1::GFP constructs used to investigate the functional relevance of FLOR-1 relocalisation upon exposure to *M. humicola* extract. The FLOR-1(79–466)::GFP, comprising the pseudokinase domain and lacking the signal peptide (SP) and transmembrane domain (TM), was designed to achieve cytosolic localisation. The FLOR-1::GFP(CAAX) construct comprises a stretch of negatively charged amino acids as well as a CAAX motif at the C-terminus for enhanced membrane association. **(B)** Representative images of the localisation of the FLOR-1::GFP constructs anterior in the head of day 1 adults as indicated. As expected, FLOR-1 kinase domain localises predominantly to the cytosol. **(C)** Representative images of *chil-27p*::GFP response to extract in FLOR-1 wild-type and basal activation of *chil-27p*::GFP in FLOR-1(79–466)::GFP without extract (see zoomed in images), which is not increased by continuous exposure to *M. humicola* extract for 72 h. **(D)** qPCR analysis of *Y71H2AR.2* and *B0507.8* shows enhanced transcript levels in FLOR-1(79-466)::GFP untreated (-) compared to untreated N2 control (-) and extract treated N2 (+) confirming the basal activity observed in (c) also for other ORR genes (biological triplicates). **(E)** Representative images of *chil-27p*::GFP response to continuous extract exposure for 72 h in wild-type, FLOR-1::GFP and FLOR-1::GFP(CAAX). **(F)** Quantification of *chil-27p*::GFP response after continuous extract exposure of wild-type (FLOR-1), FLOR-1::GFP and FLOR-1::GFP(CAAX) as shown in (E), (n=30 per condition, biological triplicates). **(G)** Model for the proposed activation and relocalisation of FLOR-1 upon extract exposure by the CDC-37/HSP-90 complex. Upon pathogen extract exposure the membrane-associated pseudokinase FLOR-1 is triggered and relocalises with support of the CDC-37/HSP-90 complex to the cytosol where it facilitates the induction of the ORR. Following activation of gene expression FLOR-1 shuttles back to the membrane for further signal perception. In contrast, the active protein kinase OLD-1 functions independently of the CDC-37/HSP-90 complex. The induction of ORR in response to *M. humicola* extract as well as the induction by OLD-1 overexpression are dependent on the membrane associated FLOR-1. Scale bars: (B) = 20 μM, (C, E) = 200 μM and 20 μM for zoom. Bars in (D) and (F) are mean±SD. (D, F) = \*\*\**p* < 0.001, \**p* < 0.05 unpaired t-test.

Overexpression of OLD-1 is sufficient to activate *chil-27p*::GFP in the absence of extract treatment (*9*). However, with cytosolic FLOR-1(79-466)::GFP the constitutive *chil-27p*::GFP activation driven by OLD-1 overexpression was abolished, and could not be restored upon supply of *M. humicola* extract (Fig. S4B). For comparison to FLOR-1(79-466)::GFP containing only the pseudokinase domain, a cytosolic OLD-1::GFP transgene similarly lacking the signal peptide and transmembrane helix (1-102 amino acids) was generated (Fig. S4C). Surprisingly while overexpression of full-length OLD-1 induced *chil-27p*::GFP without extract treatment, cytosolic OLD-1(103-522)::GFP was not sufficient to activate *chil- 27p*::GFP, and showed reduced endogenous *chil-27* transcript levels (Fig. S4D and S4E).

The basal activation by FLOR-1(79-466)::GFP in the absence of extract supports a model of FLOR-1 also having a role in the cytosol following subcellular relocalisation. Hence, the *chil-27p*::GFP response to extract as well as induction by OLD-1 overexpression was investigated upon introducing a C-terminal prenylation motif (CAAX motif) with a short stretch of negatively charged amino acids with genome editing into FLOR-1 (Fig. 4A) to further anchor FLOR-1 to the membrane. As expected, FLOR-1::GFP(CAAX) localised similarly to the wild-type protein (Fig. 4B), but extract response (Fig. 4E and 4F) as well as induction by OLD-1 overexpression (Fig. S4F and S4G) were both significantly reduced when FLOR-1 is trapped at the membrane. Taken together, these results suggest that shuttling of FLOR-1 between the membrane and the cytosol is required for complete response to *M. humicola* extract.

## DISCUSSION

Our study highlights the translational regulation of the immune response of *C. elegans* to an infection with *M. humicola* by the kinase-specific co-chaperone CDC-37 stabilising the receptor-like pseudokinase FLOR-1. TMT proteomics of the membrane-associated receptor-like pseudokinase FLOR-1 upon extract exposure identified proteins involved in signalling, proteolysis and trafficking pathways, suggesting that FLOR-1 might occupy a key position between membrane sensing and cytosolic downstream signalling. This study focuses on the newly reported interaction between FLOR-1 and the co-chaperone CDC-37 that is known to stabilise kinases by recruiting them to the HSP90 chaperone machinery (*28*). Since FLOR-1 is lacking most residues/motifs that are essential for kinase activity(*9*), it is unlikely that CDC-37 contributes to the activation of an inherently inactive pseudokinase. Instead, CDC-37-dependent FLOR-1 localisation at the membrane appears to allow its activation following pathogen recognition, which triggers FLOR-1 relocalisation to the cytosol where CDC-37/HSP90 facilitates correct FLOR-1 folding and stabilisation, possibly promoting the formation of new protein complexes involved in signal transduction (*29–31*) (Fig. 4G). While due to technical limitations unambiguously demonstrating FLOR-1 relocalisation to the cytosol remains difficult at this point, the proposed model is consistent with the reduction of FLOR-1 membrane levels without protein degradation upon pathogen extract exposure, as well as the basal activation of the ORR by cytosolic FLOR-1. Hence, FLOR-1 has potentially evolved from a receptor scaffold into a specialised chaperone-dependent pseudokinase sensor, which shuttles between the membrane required for activation upon pathogen recognition and the cytosol for signal transduction, both with the assistance of CDC-37. How exactly FLOR-1 is triggered by pathogen extract and how it is extracted from the membrane remain currently unclear. Due to its size, the ternary FLOR-1/CDC-37/HSP-90 complex is unlikely to form directly at the membrane and full maturation is likely to occur in the cytsosol to initiate downstream signalling. Additional protein interactors participating in the signalling cascade of FLOR-1 are likely to be included in the proteomics list and warrant further investigation in the future. A common temporary component of the CDC-37/HSP- 90/client kinase complex is the abundantly expressed Ser/Thr phosphatase PPH-5, which is essential for completing the cycle by restoring the initial conformation of HSP-90 as well as the basal state of the client kinase (*21*). However, for FLOR-1 regulation the PPH-5 function appears to be dispensable, potentially because FLOR- 1 as a pseudokinase is innately inactive. Alternatively, some other phosphatase may compensate for the activity of PPH-5 resetting the CDC-37/HSP-90 platform following FLOR-1 release and activation.

Surprisingly, we found that the CDC-37/HSP-90-mediated control of the oomycete-recognition response is specifically through the pseudokinase and independent of the protein tyrosine kinase OLD-1, which is involved in the same pathway and belongs to the same KIN-16 family (*10, 32*). Although OLD-1 levels depend on FLOR-1, no FLOR-1/OLD-1 interaction was confirmed, indicating that FLOR-1 and OLD-1 do not engage in a typical kinase-pseudokinase pair (*9, 33*). Pseudokinases often retain the overall kinase fold structure and may require chaperone assistance for proper folding and maintainance of scaffold conformations (*29*). Recently, the structure of a human membrane receptor guanylyl cylase, comprising an intracellular pseudokinase domain, in complex with CDC37/HSP90 has been reported (*34*). This study provides structural evidence that pseudokinases like FLOR-1 can in fact interact with CDC-37 and that this regulatory interaction can be hijacked by proteins with functions beyond typical kinases.

The CDC37/HSP90 chaperone complex has been shown to be essential for the stability and activation of numerous protein kinases, mostly cytosolic kinases, such as AKT (Protein Kinase B), a serine/threonine kinase central to the PI3K/AKT signalling pathway thereby regulating cell survival, growth, and proliferation. However, in this case the CDC37/HSP90 complex holds the kinase in its active state in the cytosol and is not responsible for membrane recruitment and subsequent activation (*35, 36*) Similarly, also RAF, the cytosolic Ser/Thr kinase of the MAPK/ERK cascade which is activated by membrane-bound RAS, is stabilised by CDC37/HSP90 in its active form (*23, 37, 38, 39*). While primarily acting intracellularly, the CDC-37/HSP-90 complex has also been shown to stabilise membrane-associated tyrosine kinase HER2 of the ERBB family (*14*). HER2/ERBB2 stability is maintained through HSP90 chaperoning, and this interaction also occurs at the cell surface, primarily in cancer cells (*40, 41*).

HSP-90 has been previously shown to influence immune responses in *C. elegans* (*42*), and this work provides a mechanistic link between the CDC-37/HSP- 90 complex and oomycete recognition. Furthermore, our study illustrates that the function of CDC-37 in the case of FLOR-1 goes beyond classic kinase activation and likely involves dynamic relocalisation of the pseudokinase from the membrane to the cytosol, comparable to the functional relocalisation of other kinases to the membrane or nucleus (*43, 44*).

The wide involvement of CDC-37 in kinase quality control and cellular signalling underscores its importance in various processes such as cell cycle control, development, differentiation and stress response (*36, 38, 45, 46*) as well as diseases ranging from Parkinson and Alzheimer’s to cancer (*47–49*). In mammalian systems, CDC37 is commonly implicated in oncogenic processes by stabilising oncogenic kinases, facilitating signalling networks and promoting proliferative phenotypes during tumorigenesis. Hence, it is not surprising that current strategies also focus on disrupting the CDC-37/HSP-90 interaction to selectively destabilise oncogenic kinases and prevent tumour growth and metastasis (*50*). Improving our understanding of the regulation and functions of pseudokinases in physiological as well as pathological settings can provide valuable insights into their roles in cellular signalling and new opportunities for therapeutic intervention (*51, 52*).

## MATERIAL AND METHODS

### *C. elegans* strain maintenance

All *C. elegans* strains are maintained under standard conditions on NGM plates seeded with *E. coli* OP50 at 20 °C. The strains used in this study are listed in Table S5.

### Molecular cloning and transgenesis

To obtain a plasmid coding for *cdc-37p*::CDC-37::mScarlet the *cdc-37* locus including an additional 8 bp upstream until the next coding sequence was amplified from gDNA (N2) using primers FT150/FT151. The vector backbone pSEM318 including a C-terminal mScarlet was amplified with FT152/FT153. Sequences of oligos are listed in Table S6. Both fragments were ligated with a Gibson reaction creating pFT39 which was verified by full plasmid sequencing (Full Circle Labs). pFT39 was injected into FLOR-1::GFP (MBA998) at 25 ng/μl together with a hygromycin resistance marker (pCFJ782) at 15 ng/μl and pBJ36 as carrier DNA to a total concentration of 110 ng/μl in the mix. To obtain a plasmid coding for dpy-7::old- 1(103–522)::GFP the old-1 locus was amplified from pFT43 using primers FT216/FT217. The vector backbone pIR6 including an N-terminal dpy-7 promotor was amplified with FT218/FT219. Both fragments were ligated with a Gibson reaction creating pFT44 which was verified by full plasmid sequencing (Full Circle Labs).

### CRISPR-mediated genome editing

To generate a strain containing the cytosolic, kinase domain only FLOR-1(79–466)::GFP, crRNA 1 and 2 were used with one binding in the 5‘ UTR and the other one inside *flor-1* to generate *flor-1(icb221)*. The sequences of crRNA used in this study can be found in Table S6. For the injection mix, 1.4 µl of each crRNA (34 µM stock, IDT) was added to 5 µl of tracrRNA (18 µM stock, IDT) and 1 µl of Cas9 protein (2.5 µg/µl stock, IDT) an mixed by pipetting before incubating at 37 °C for 15 min. After incubation, 2.2 μl of a single stranded donor template (25 μM stock, IDT), 5 ng/μl of myo-2::GFP and water to a final volume of 20 μl was added. The mix was centrifuged at 20,000 xg for 5 min and used to inject MBA998 (FLOR-1::GFP). myo-2::GFP positive F1 were singled out and screened for heterozygosity with FT172/FT173. F2 were singled out and genotyped for homozygous deletion with T01G5.1 gene F swaI site/FT173. Successful modification was confirmed by Sanger sequencing.

The same procedure was followed to create FLOR-1::GFP(CAAX) by adding a CAAX motif containing a short stretch of negatively charged amino acids C-terminal of *flor-1::gfp*. crRNA 3 with a binding site inside GFP of *flor-1::gfp* and the respective donor template were used to generate *flor-1(icb223)*. myo-2::GFP positive F1 were singled out and screened for heterozygosity with FT220/FT221. F3 was checked for homozygosity with FT220/FT221. Successful integration of the CAAX motif was confirmed by Sanger sequencing.

### Extract treatment

Oomycete extracts from *M. humicola* were produced as previously described (*8*). Extract treatment was performed by adding 300 μl of extract onto NGM plates seeded with OP50. For continuous extract treatment eggs of day 1 gravid adults were plated onto extract covered plates and day 1 adults imaged and scored. For extract response after RNAi treatment, extract was added at L4 stage of the following generation and animals were imaged as well as scored after 24 h.

### RNA Interference (RNAi)

RNAi feeing was performed using *E. coli* HT115 expressing double-stranded RNA corresponding to the target gene or the empty vector as a control. NGM plates supplemented with 25 μg/ml ampicillin, 6.25 μg/ml tetracyclin and 1 mM IPTG, were seeded with 4x concentrated bacterial culture. All RNAi treatments started at L4 stage and day 1 adults of the next generation were imaged and scored. Apart from the initial RNAi screen (Table S4), all RNAi treatments were performed in biological triplicates. RNAi clones used in this study were obtained from the Ahringer RNAi Library (Source Bioscience, (*53*)) and verified by Sanger Sequencing. The *hsp-90* and *pph-5* RNAi clones were obtained from the ORFeome Library (*54*). To circumvent larval arrest, *hsp-90* RNAi was diluted 1:1 with control RNAi.

### Microscopy

For imaging animals are picked into a drop of M9 with 50 μM sodium azide on a 3 % agarose pad mounted onto a glas slide. To document the *chil-27p*::GFP expression in response to extract in the initial RNAi screen, a Zeiss Axio Zoom V16 microscope was used at 112x magnification with a 38 HE GFP fluorescence filter (470 ± 40/ 525 ± 50 nm). For all other conditions confocal microscopy was performed with a Leica SP8-Stellaris 5 Inverted Light Sheet confocal microscope (GFP: 485/ 490-577 nm, DsRed: 555/ 560-734 nm, mScarlet: 569/ 574-700 nm) with a HC PL APO 10x/0.40 CS2 or HC PL APO CS2 63x/1.40 Oil objective. FLOR-1 and OLD-1 levels were measured as total GFP fluorescence intensity per area of the head. To measure relocalisation of FLOR-1 to the cytosol, the average of 3 random areas in the head area away from membrane structures was compared to the total amount of protein detected in the whole head and the ratio is presented.

### RNA extraction, cDNA synthesis and qPCR

RNA extraction was performed with three biological replicates per condition. Pellets of *C. elegans* were resuspended in TRIzol™ (Invitrgen) in a ratio 1:5 (v/v) and animals lysed by repeated snap freezing in liquid nitrogen and reheating at 37 °C with subsequent vortexing. Next, chloroform (15 % of the initial TRIzol™ volume) was added and layers separated by centrifugation at 20,000 xg for 15 min at 4 °C. The top phase was transfered to isopropanol (50 % of initial TRIzol™ volume), mixed and incubated for 10 min at RT. The RNA was pelleted by centrifugation at 20,000 xg for 15 min at 4 °C. The supernatant was removed and the pellet washed with 500 μl of 70 % ethanol. The RNA pellet was collected at 20,000 xg for 5 min at RT and air-dried before being resuspended and used for cDNA synthesis.

cDNA was prepared using Superscript IV VILO Master mix (Thermo Fisher Scientific) according to manufacturer’s instructions with some minor modifications. Incubation at 37 °C and at 50 °C were extended to 15 min and 30 min, respectively. qPCR was performed with SYBR™ Select Master Mix for CFX (Applied Biosystems) according to manufacturer’s instructions on a CFX Connect Real-Time PCR Detection System (Bio-Rad). In total 45 cycles were run with a 1 min annealing/extension step at 60 °C. Each condition is normalised to *rpl-26* and N2 with no treatment (N2-) resulting in ΔΔCt values. The sequences of qPCR primers used can be found in Table S6.

### co-Immunoprecipitation in *C. elegans*

For protein extraction *C. elegans* populations expressing proteins as indicated were scaled up to approximately 2x10^6^ of synchronised animals on 8P plates (3 g/l NaCl, 20 g/l bactopeptone, 25 g/l agar). Animals were collected in M9 and pelleted at 1,000 xg for 10 min. The supernatant was discarded and the worm suspension dropped into a mortar filled with liquid nitrogen for grinding. After grinding for 10 min the powder was resuspended in 10 ml extraction buffer (25 mM Tris, pH 8.0, 1 mM DTT (Dithiothreitol), 150 mM NaCl, 1 mM PMSF (phenylmethylsulfonyl fluoride), protease inhibitor cocktail (Pierce), Phosphostop^TM^ (Roche) and 0.2 % NP40) and subsequently, sonicated 10 times for 30 sec with 1 min interval at 50% (Sonics, Vibra-Cell™ VCX 130). The suspension was then centrifuged for 30 min at 20,000 xg and the cleared supernatant was subjected to 20 ul of equelibrated GFP or RFP agarose beads (proteintech). Beads were incubated for 1.5 h on a rotator at 4 °C. Finally, beads were washed 3 times with extraction buffer before proteins were eluted at 70 °C for 10 min in 35 ul SDS loading buffer.

### Immuno blot

Samples were run on a NuPAGE™ 4 to 12%, Bis-Tris Mini gel and then transferred at 30 V for 70 min to a nitrocellulose membrane using a Mini gel tank (Invitrogen). Membranes were blocked in 3 % BSA in PBST (137 mM NaCl, 12 mM phosphate, 2.7 mM KCl, pH 7.4, 0.1 % tween-20) before adding the primary antibody at a 1:1500 dilution (α-GFP, rabbit, Invitrogen, A11122; α-mCherry, rabbit, Abcam, ab167453) for incubation overnight at 4 °C on a shaker. The membrane was washed 4 times with PBST before adding the secondary HRP-coupled anti-rabbit antibody at 1:3000 dilution for 1 h at RT (Sigma, A9169). After another 4 washing steps with PBST, blots were visualised with ECL (Amersham) on a Odyssey Fc (Licor) or ChemiDoc MP Imaging System (BioRad).

### TMT Labelling and high pH reversed-phase chromatography

For TMT proteomics each condition was performed in biological triplicates. Protein aggregates in IP eluates were removed by running samples into the separating gel of a 10 % SDS-Page. Lanes were excised and subjected to in-gel tryptic digestion using a DigestPro automated digestion unit (Intavis). The peptides were evaporated, resuspended in 50 ul of 100 mM Triethylammonium bicarbonate buffer and labelled with TMTpro 16plex Label Reagent according to the manufacturer’s protocol (Thermo Fisher Scientific). The samples were pooled and desalted using a SepPak cartridge according to the manufacturer’s instructions (Waters). The eluate from the SepPak cartridge was evaporated and resuspended in buffer A (20 mM ammonium hydroxide, pH 10) prior to fractionation by high pH reversed-phase chromatography using an Ultimate 3000 liquid chromatography system (Thermo Fisher Scientific). The sample was loaded onto a XBridge BEH C18 Column (Waters) in buffer A and peptides eluted with an increasing gradient of buffer B (20 mM Ammonium Hydroxide in acetonitrile, pH 10) from 0-95 % over 1 h. The 6 resulting fractions were evaporated and resuspended in 1 % formic acid

### Nano-LC Mass Spectrometry

High pH reversed-phase fractions were further fractionated using an Ultimate 3000 nano-LC system in line with an Orbitrap Fusion Lumos mass spectrometer (Thermo Scientific). Peptides in 1 % formic acid were loaded onto an Acclaim PepMap C18 nano-trap column (Thermo Scientific). After washing with 0.5 % acetonitrile, 0.1 % formic acid peptides were resolved on a 250 mm × 75 μm Acclaim PepMap C18 reverse phase analytical column (Thermo Scientific) over a 2.5 h organic gradient. Peptides were ionised by nano-electrospray ionisation at 2 kV using a stainless-steel emitter at 300 °C. All spectra were acquired using an Orbitrap Fusion Lumos mass spectrometer controlled by Xcalibur 3.0 software (Thermo Scientific) and operated in data-dependent acquisition mode using an SPS-MS3 workflow.

### Data analysis of Nano-LC Mass Spectra

The raw data files were processed and quantified using Proteome Discoverer software v2.4 (Thermo Scientific) and searched against the UniProt *C. elegans* database (September 2023: 26688 entries) using the SEQUEST HT algorithm. Peptide precursor mass tolerance was set at 10 ppm and MS/MS tolerance at 0.6 Da. Search criteria included oxidation of methionine, acetylation of the N-terminus, carbamidomethylation of cysteine and the addition of the TMTpro 16plex tag (+304.207 Da) to peptide N-termini and lysines as fixed modifications. Searches were performed with full tryptic digestion and a maximum of 2 missed cleavages were allowed. The reverse database search option was enabled and all data was filtered to satisfy a false discovery rate (FDR) of 5 %.

### Proteomics analysis with Perseus

TMT scores of all samples were normalised to the sample with the highest GFP::CAAX score and read in to Perseus 2.0.11.0. After a log2(x) transformation, missing values were replaced from normal distribution and the average subtracted in order to reduce noise. In total 983 hits were used for principal component analysis and for the determination of significance with a permutation-based FDR value of 0.05 with a t-test.

### Statistics Quantification and statistical analysis

Graphic representation and statistical analysis were performed using GraphPad Prism 7. Data shown in bar graphs indicate mean, and error bars standard deviation. An unpaired t-test was used to evaluate significance. Results were considered statistically significant when p < 0.05. Asterisks in figures indicate corresponding statistical significance as follows: ∗p < 0.05; ∗∗∗p < 0.001, n.s stands for not significant.

## ACKNOWLEDGMENTS

We thank Florence Drury, Ming Yi and Jonathan Saunders for strains. Some strains are provided by the Caenorhabditis Genetics Center (CGC), which is funded by NIH Office of Research Infrastructure Programs (P40 OD010440). We also thank the Facility for Imaging by Light Microscopy (FILM) at Imperial College London, which is partly supported by funding from the BBSRC (BB/T017929/1).

## STATEMENTS AND DECLARATIONS

Funding

This work was supported by the BBSRC (BB/X001865/1).

## Contributions

F.T. conducted experiments and analysed the data together with M.B. F.T. and M.B. wrote the manuscript.

## Data availability

All strains and oligos used in this study are described in Tables S5-6 and are available upon request. All data necessary for confirming the conclusions of the work are present within the article and figures. Supplemental material is available online.

**Table S1: Normalised TMT data of FLOR-1::GFP compared to GFP::CAAX.**

**Table S2: Normalised TMT data of FLOR-1::GFP with and without extract.**

**Table S3: Normalised TMT data of OLD-1::GFP compared to GFP::CAAX.**

**Table S4: Selected candidates from TMT data used for the RNAi screen.**

**Table S5: Strains used in this study.**

**Table S6: Primers and oligos used in this study.**

---

**Figure S1.**
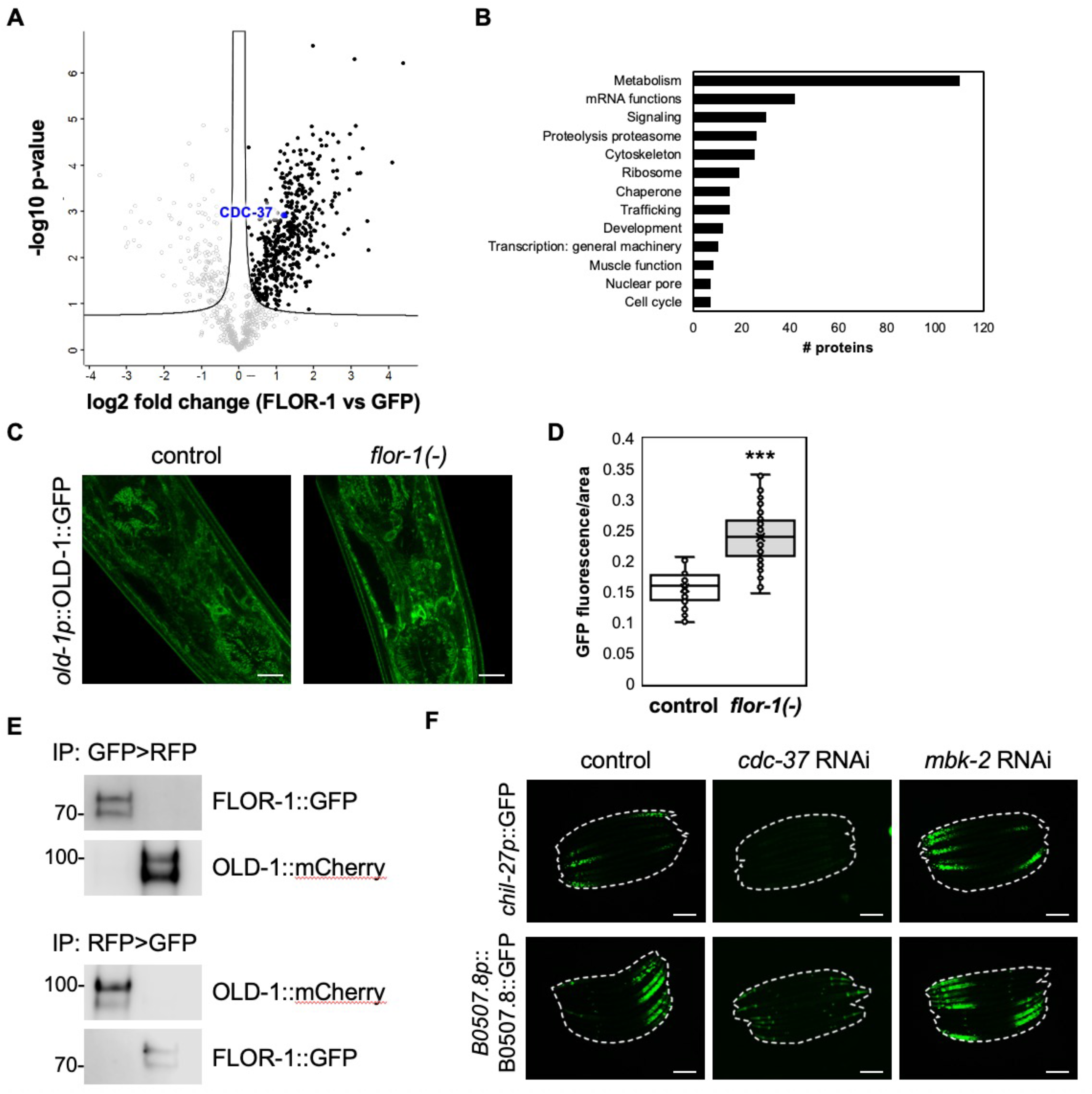
TMT-based proteomic analysis identifies the kinase-specific co-chaperone CDC-37 as an ORR regulator. **(A)** Volcano plot visualising log_2_ fold change and statistical significance of FLOR-1::GFP interactors compared to GFP::CAAX. A total of 479 proteins showed significant differential enrichment (FDR < 0.05, filled circles) amongst 983 identified proteins in total. The co-chaperone CDC-37 studied here is highlighted in blue. **(B)** Functional gene set enrichment analysis using WormCat 2.0 of significantly enriched FLOR-1::GFP interactors compared to GFP::CAAX (p < 0.05) identifies categories amongst others associated with signalling, proteolysis and trafficking. **(C)** Representative images showing increased membrane localization of *old-1p*::OLD-1::GFP in the anterior head region of day 1 adults in *flor-1(icb115)* compared to wild-type. **(D)** Quantification of OLD-1::GFP fluorescence per area anterior in the head of day 1 adults as shown in (C), (n=40 per condition). **(E)** Co-immunoprecipitation in animals expressing FLOR-1::GFP and *dpy-7p*::OLD-1::mCherry using either GFP- or RFP-trapping confirm the lack of an interaction as anticipated from TMT proteomics. Reciprocal co-immunoprecipitations have been performed to verify enough protein in each sample. Molecular weights are indicated in kilodaltons (kDa). **(F)** Representative images of *chil-27p*::GFP and *B0507.8p*::B0507.8::GFP induction upon 24 h *M. humicola* extract exposure following control, *cdc-37*, and *mbk-2* RNAi. Whereas *cdc-37* RNAi completely abolishes extract response in both reporters, RNAi of the related dual specificity kinase *mbk-2* has no effect. Scale bars: (C) = 10 μM, (F) = 200 μM. (D) = \*\*\**p* < 0.001, unpaired t-test.

**Figure S2.**
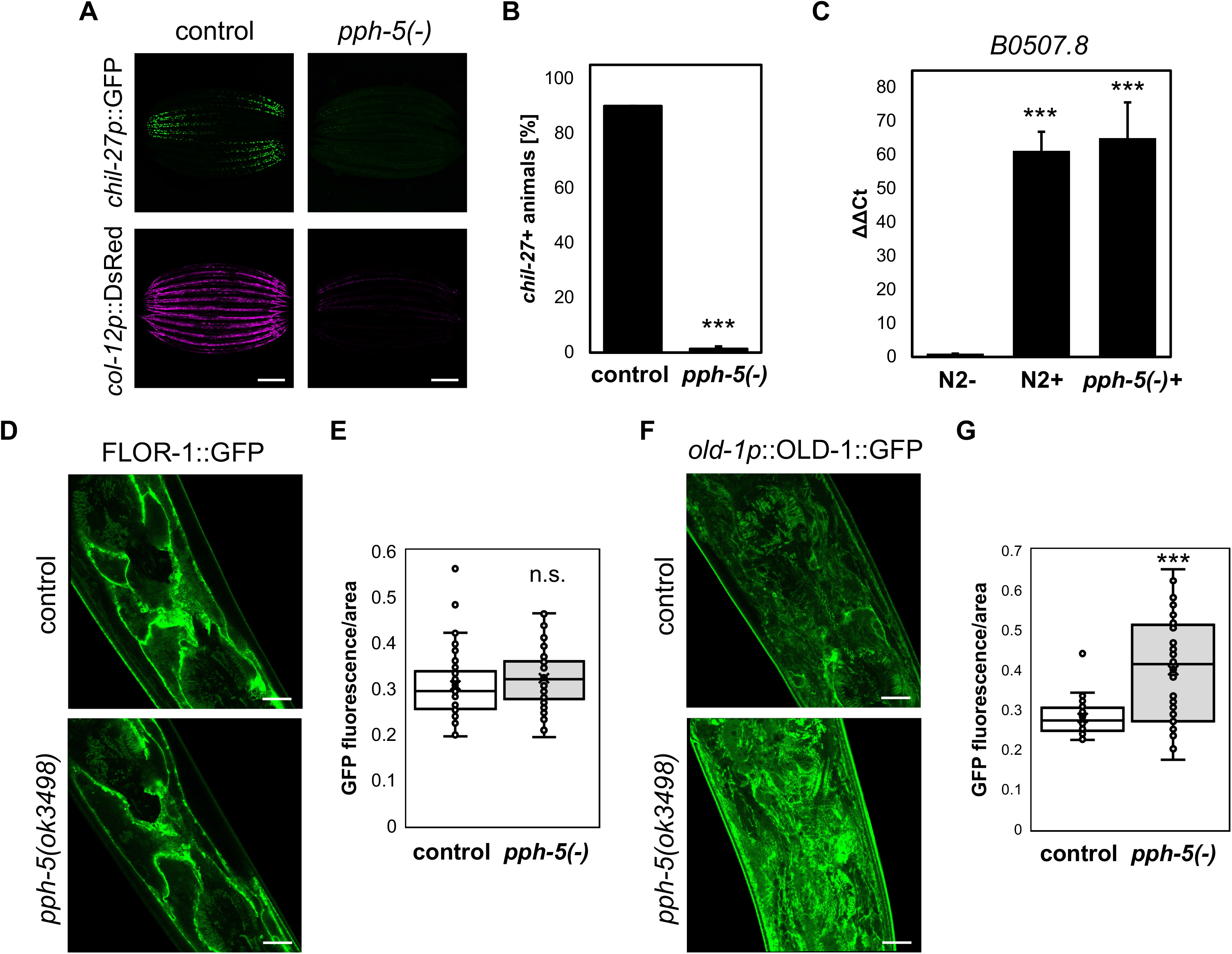
The CDC-37/HSP-90 complex-specific phosphatase PPH-5 is dispensable for the chil-27p::GFP response to M. humicola extract. **(A)** Representative images of *chil-27p*::GFP expression in a *pph-5(ok3498)* deletion mutant after 72 h *M. humicola* extract exposure. The *chil-27p*::GFP response but also *col-12p*::DsRed expression are significantly reduced. **(B)** Quantification of *chil-27p*::GFP in *pph-5(ok3498)* animals in response to extract as shown in (A), (n=30 per condition, biological triplicates). **(C)** qPCR analysis of B0507.8 upon extract exposure for 72 h (+) of animals in *pph-5* wildtype and deletion mutants, normalised to untreated N2 control (-), suggests the *chil-27p*::GFP response in (A) arises from transgene silencing rather than impaired ORR activation. (biological triplicates). **(D)** Representative images showing FLOR-1::GFP localisation anterior in the head of day 1 adults. No differences of FLOR-1::GFP membrane localisation are observed between wild-type and *pph-5(ok3498)*. **(E)** Quantification of FLOR-1::GFP intensity per area anterior in the head of day 1 adults as shown in (D), (n=40 per condition). **(F)** Representative images of the localisation of OLD-1::GFP anterior in the head of day 1 adults indicate that in *pph-5(ok3498)* mutants, OLD-1::GFP accumulates at the membrane. **(G)** Quantification of OLD-1::GFP intensity per area anterior in the head of day 1 adults in *pph-5(ok3498)* as shown in (F), (n=40 per condition). Scale bars: (A) = 200 μM, (D, F) = 10 μM. Bars in (B) and (C) are mean±SD. (B, C, E, G) = \*\*\**p* < 0.001, unpaired t-test.

**Figure S3.**
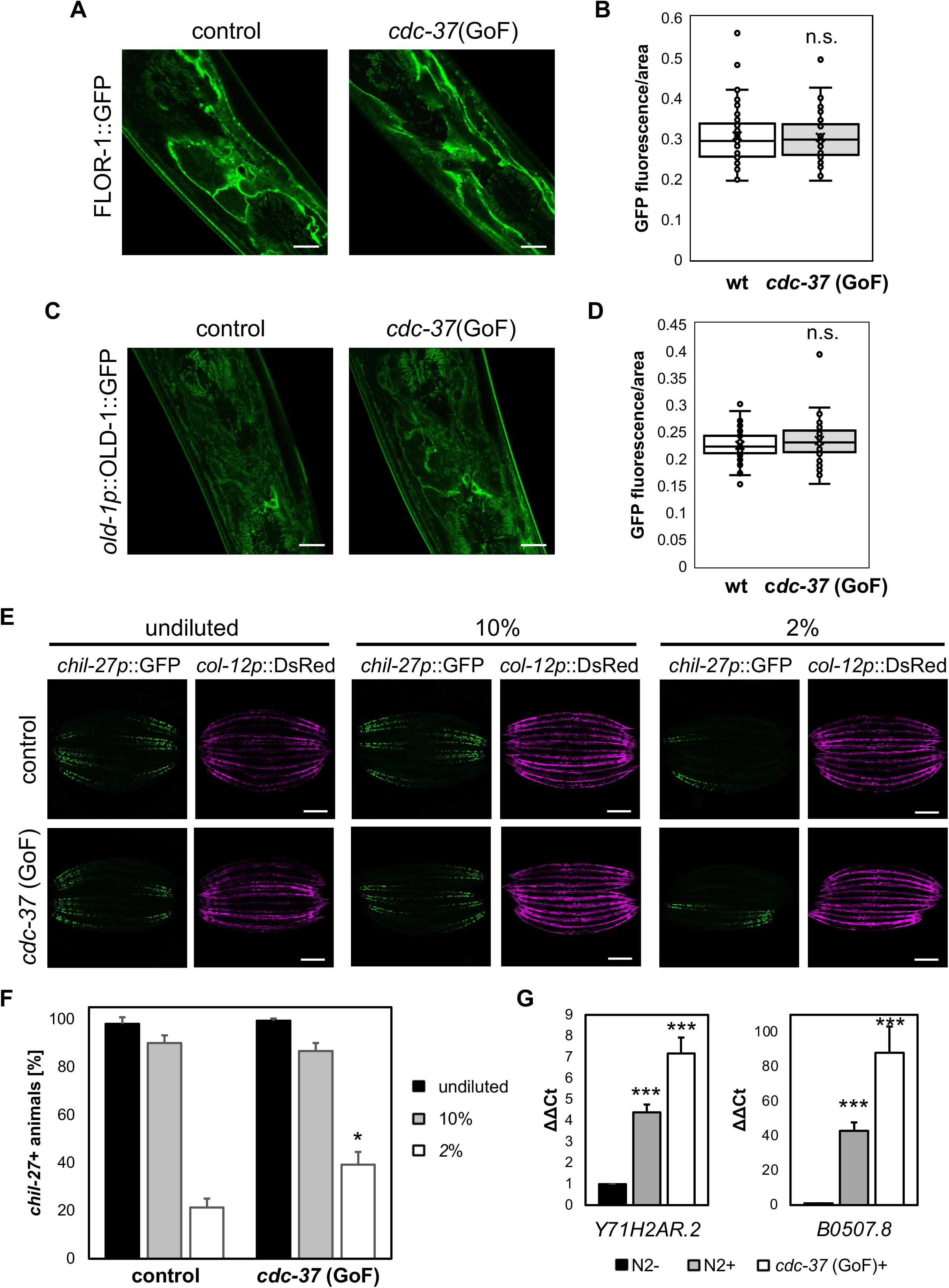
A cdc-37 gain-of-function allele enhances the M. humicola extract-induced ORR response. **(A)** Representative images show no difference in FLOR-1::GFP localisation anterior in the head of day 1 adults of wild-type and *cdc-37(ax2001)* gain-of-function (GoF) animals. **(B)** Quantification of FLOR-1::GFP intensity per area anterior in the head of day 1 adults as show in (A), (n=40 per condition). **(C)** Representative images of membrane-associated *old-1p*::OLD-1::GFP anterior in the head of day 1 adults of wild-type and *cdc-37(ax2001)* GoF animals. **(D)** Quantification of OLD-1::GFP intensity per area anterior in the head of day 1 adults as shown in (C). **(E)** Representative images of c*hil-27p*::GFP expression in wild-type and a *cdc-37(ax2001)* GoF mutant in response to continuous exposure to dilutions of *M. humicola* extract for 72 h. At higher extract dilutions *cdc-37(ax2011)* GoF significantly enhances *chil-27p*::GFP response. **(F)** Quantification of *chil-27p*::GFP response after continuous extract exposure at different dilutions of wild-type and cdc-37*(ax2001)* GoF animals as shown in (E), (n=30 per condition, biological triplicates). **(G)** qPCR analysis shows elevated transcript levels of *Y71H2AR.2* and *B0507.8* in *cdc-37(ax2001)* GoF after continuous extract exposure for 72 h (*cdc-37* GoF+) in comparison to N2 treated with extract (+), normalised to N2 without extract treatment (-). Scale bars: (A, C) = 10 μM, (E) = 200 μM. Bars in (F) and (G) are mean±SD. (F) = \**p* < 0.05, (G) = \*\*\**p* < 0.001, unpaired t-test.

**Figure S4.**
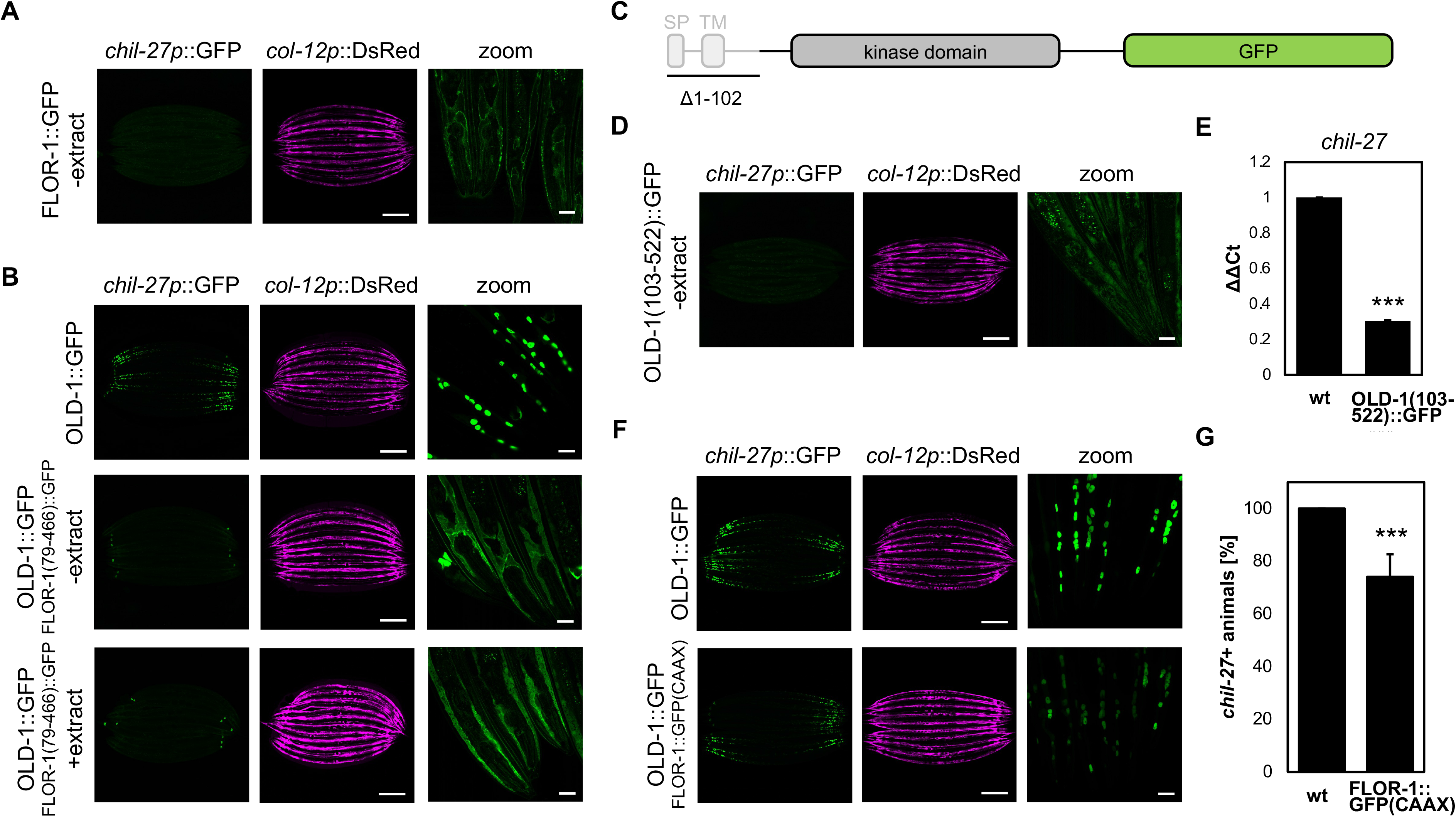
**The activation of *chil-27p*::GFP by OLD-1 overexpression is dependent on accurate FLOR-1 localisation**. **(A)** Representative images show the lack of basal activation of *chil-27p*::GFP in FLOR-1 wild-type without extract. **(B)** Representative images of *chil-27p*::GFP induction by OLD-1 overexpression in wild-type and the lack of *chil-27p*::GFP induction in FLOR-1(79-466)::GFP, which is also not restored after continuous exposure to to *M. humicola* extract for 72 h. **(C)** Schematic diagram of *dpy-7*::OLD-1(103-522)::GFP, comprising the kinase domain and lacking the signal peptide (SP) and transmembrane domain (TM). **(D)** Representative images show the lack of basal activation of *chil-27p*::GFP in OLD-1(103-522)::GFP in the absence of extract, in contrast to FLOR-1(79-466)::GFP. **(E)** qPCR analysis of *chil-27p*::GFP shows reduced transcript levels in OLD-1(103-522)::GFP compared to untreated N2 control and confirms the lack of basal activity observed in (C), (biological triplicates). **(F)** Representative images of *chil-27p*::GFP induction by OLD-1 overexpression in FLOR-1::GFP(CAAX) is reduced compared to wild-type. **(G)** Quantification of *chil-27p*::GFP induction by OLD-1 overexpression in wild-type and FLOR-1::GFP(CAAX) as shown in (F), (n=30 per condition, biological triplicates). Scale bars: (A, C, E, F) = 200 μM and 20 μM for zoom. Bars in (D) and (G) are mean±SD. (D, G) = \*\*\**p* < 0.001, unpaired t-test.

## REFERENCES

1. H. Schulenburg, M. A. Felix, The Natural Biotic Environment of *Caenorhabditis elegans*. Genetics 206, 55–86 (2017).

2. S. Tse-Kang, K. A. Wani, R. Pukkila-Worley, Patterns of pathogenesis in innate immunity: insights from *C. elegans*. Nat Rev Immunol, (2025).

3. G. A. Osman et al., Natural Infection of *C. elegans* by an Oomycete Reveals a New Pathogen-Specific Immune Response. Curr Biol 28, 640–648 e645 (2018).

4. L. Derevnina et al., Emerging oomycete threats to plants and animals. Philos Trans R Soc Lond B Biol Sci 371, (2016).

5. T. D. Tran, R. J. Luallen, An organismal understanding of *C. elegans* innate immune responses, from pathogen recognition to multigenerational resistance. Semin Cell Dev Biol 154, 77–84 (2024).

6. E. A. Evans, T. Kawli, M. W. Tan, Pseudomonas aeruginosa suppresses host immunity by activating the DAF-2 insulin-like signaling pathway in *Caenorhabditis elegans*. PLoS Pathog 4, e1000175 (2008).

7. T. Kitisin, W. Muangkaew, P. Sukphopetch, *Caenorhabditis elegans* DAF-16 regulates lifespan and immune responses to Cryptococcus neoformans and Cryptococcus gattii infections. BMC Microbiol 22, 162 (2022).

8. M. K. Fasseas et al., Chemosensory Neurons Modulate the Response to Oomycete Recognition in *Caenorhabditis elegans*. Cell Rep 34, 108604 (2021).

9. F. Drury et al., A PAX6-regulated receptor tyrosine kinase pairs with a pseudokinase to activate immune defense upon oomycete recognition in *Caenorhabditis elegans*. Proc Natl Acad Sci U S A 120, e2300587120 (2023).

10. B. A. Rikke, S. Murakami, T. E. Johnson, Paralogy and orthology of tyrosine kinases that can extend the life span of *Caenorhabditis elegans*. Mol Biol Evol 17, 671–683 (2000).

11. G. Manning, Genomic overview of protein kinases. WormBook, 1–19 (2005).

12. M. Taipale et al., Quantitative analysis of HSP90-client interactions reveals principles of substrate recognition. Cell 150, 987–1001 (2012).

13. D. Keramisanou et al., Molecular Mechanism of Protein Kinase Recognition and Sorting by the Hsp90 Kinome-Specific Cochaperone Cdc37. Mol Cell 62, 260–271 (2016).

14. J. R. Smith, P. A. Clarke, E. de Billy, P. Workman, Silencing the cochaperone CDC37 destabilizes kinase clients and sensitizes cancer cells to HSP90 inhibitors. Oncogene 28, 157–169 (2009).

15. A. P. Kornev, S. S. Taylor, Dynamics-Driven Allostery in Protein Kinases. Trends Biochem Sci 40, 628–647 (2015).

16. K. A. Verba et al., Atomic structure of Hsp90-Cdc37-Cdk4 reveals that Hsp90 traps and stabilizes an unfolded kinase. Science 352, 1542–1547 (2016).

17. J. Shao, A. Irwin, S. D. Hartson, R. L. Matts, Functional dissection of cdc37: characterization of domain structure and amino acid residues critical for protein kinase binding. Biochemistry 42, 12577–12588 (2003).

18. S. M. Roe et al., The Mechanism of Hsp90 regulation by the protein kinase-specific cochaperone p50(cdc37). Cell 116, 87–98 (2004).

19. J. M. Eckl et al., Cdc37 (cell division cycle 37) restricts Hsp90 (heat shock protein 90) motility by interaction with N-terminal and middle domain binding sites. J Biol Chem 288, 16032–16042 (2013).

20. J. M. Eckl et al., Hsp90.Cdc37 Complexes with Protein Kinases Form Cooperatively with Multiple Distinct Interaction Sites. J Biol Chem 290, 30843–30854 (2015).

21. M. Jaime-Garza et al., Hsp90 provides a platform for kinase dephosphorylation by PP5. Nat Commun 14, 2197 (2023).

22. G. Chen, P. Cao, D. V. Goeddel, TNF-induced recruitment and activation of the IKK complex require Cdc37 and Hsp90. Mol Cell 9, 401–410 (2002).

23. S. Garcia-Alonso et al., Structure of the RAF1-HSP90-CDC37 complex reveals the basis of RAF1 regulation. Mol Cell 82, 3438–3452 e3438 (2022).

24. M. L. Stitzel, J. Pellettieri, G. Seydoux, The *C. elegans* DYRK Kinase MBK-2 Marks Oocyte Proteins for Degradation in Response to Meiotic Maturation. Curr Biol 16, 56–62 (2006).

25. K. Liu et al., Paired C-type lectin receptors mediate specific recognition of divergent oomycete pathogens in *C. elegans*. Cell Rep 43, 114906 (2024).

26. J. Cao et al., Comprehensive single-cell transcriptional profiling of a multicellular organism. Science 357, 661–667 (2017).

27. Y. Wang et al., Identification of suppressors of mbk-2/DYRK by whole-genome sequencing. *G3* (*Bethesda*) 4, 231–241 (2014).

28. M. Taipale, D. F. Jarosz, S. Lindquist, HSP90 at the hub of protein homeostasis: emerging mechanistic insights. Nat Rev Mol Cell Biol 11, 515–528 (2010).

29. J. Boudeau, D. Miranda-Saavedra, G. J. Barton, D. R. Alessi, Emerging roles of pseudokinases. Trends Cell Biol 16, 443–452 (2006).

30. K. A. Verba, D. A. Agard, How Hsp90 and Cdc37 Lubricate Kinase Molecular Switches. Trends Biochem Sci 42, 799–811 (2017).

31. D. P. Byrne et al., Evolutionary and cellular analysis of the ’dark’ pseudokinase PSKH2. Biochem J 480, 141–160 (2023).

32. C. Popovici, R. Roubin, F. Coulier, P. Pontarotti, D. Birnbaum, The family of Caenorhabditis elegans tyrosine kinase receptors: similarities and differences with mammalian receptors. Genome Res 9, 1026–1039 (1999).

33. A. W. Smith, F. N. Barrera, Regulation of receptor tyrosine kinase hetero-interactions. Curr Opin Struct Biol 95, 103187 (2025).

34. N. A. Caveney, N. Tsutsumi, K. C. Garcia, Structural insight into guanylyl cyclase receptor hijacking of the kinase-Hsp90 regulatory mechanism. Elife 12, (2023).

35. A. D. Basso et al., Akt forms an intracellular complex with heat shock protein 90 (Hsp90) and Cdc37 and is destabilized by inhibitors of Hsp90 function. J Biol Chem 277, 39858–39866 (2002).

36. B. D. Manning, A. Toker, AKT/PKB Signaling: Navigating the Network. Cell 169, 381–405 (2017).

37. N. Grammatikakis, J. H. Lin, A. Grammatikakis, P. N. Tsichlis, B. H. Cochran, p50(cdc37) acting in concert with Hsp90 is required for Raf-1 function. Mol Cell Biol 19, 1661–1672 (1999).

38. G. Thiel, M. Ekici, O. G. Rossler, Regulation of cellular proliferation, differentiation and cell death by activated Raf. Cell Commun Signal 7, 8 (2009).

39. L. I. Finci et al., Structural dynamics of RAF1-HSP90-CDC37 and HSP90 complexes reveal asymmetric client interactions and key structural elements. Commun Biol 7, 260 (2024).

40. A. Citri et al., Hsp90 restrains ErbB-2/HER2 signalling by limiting heterodimer formation. EMBO Rep 5, 1165–1170 (2004).

41. A. El Hamidieh, N. Grammatikakis, E. Patsavoudi, Cell surface Cdc37 participates in extracellular HSP90 mediated cancer cell invasion. PLoS One 7, e42722 (2012).

42. J. Eckl, S. Sima, K. Marcus, C. Lindemann, K. Richter, Hsp90-downregulation influences the heat-shock response, innate immune response and onset of oocyte development in nematodes. PLoS One 12, e0186386 (2017).

43. M. Wartmann, R. J. Davis, The native structure of the activated Raf protein kinase is a membrane-bound multi-subunit complex. J Biol Chem 269, 6695–6701 (1994).

44. D. Mahony, D. A. Parry, E. Lees, Active cdk6 complexes are predominantly nuclear and represent only a minority of the cdk6 in T cells. Oncogene 16, 603–611 (1998).

45. S. Bogdan, C. Klambt, Epidermal growth factor receptor signaling. Curr Biol 11, R292–295 (2001).

46. S. J. Baker, P. I. Poulikakos, H. Y. Irie, S. Parekh, E. P. Reddy, CDK4: a master regulator of the cell cycle and its role in cancer. Genes Cancer 13, 21–45 (2022).

47. A. Weihofen, B. Ostaszewski, Y. Minami, D. J. Selkoe, Pink1 Parkinson mutations, the Cdc37/Hsp90 chaperones and Parkin all influence the maturation or subcellular distribution of Pink1. Hum Mol Genet 17, 602–616 (2008).

48. S. K. Calderwood, Cdc37 as a co-chaperone to Hsp90. Subcell Biochem 78, 103–112 (2015).

49. L. Gracia, G. Lora, L. J. Blair, U. K. Jinwal, Therapeutic Potential of the Hsp90/Cdc37 Interaction in Neurodegenerative Diseases. Front Neurosci 13, 1263 (2019).

50. L. Wang, Q. Zhang, Q. You, Targeting the HSP90-CDC37-kinase chaperone cycle: A promising therapeutic strategy for cancer. Med Res Rev 42, 156–182 (2022).

51. A. Virtanen et al., Molecular basis of JAK kinase regulation guiding therapeutic approaches: Evaluating the JAK3 pseudokinase domain as a drug target. Adv Biol Regul 95, 101072 (2025).

52. S. R. R. Gupta, R. Rameshwari, I. K. Singh, ROR1 protein: a pseudokinase at the crossroads of cancer progression and therapy. Mol Biol Rep 53, 257 (2026).

53. R. S. Kamath et al., Systematic functional analysis of the *Caenorhabditis elegans* genome using RNAi. Nature 421, 231–237 (2003).

54. J. Reboul et al., *C. elegans* ORFeome version 1.1: experimental verification of the genome annotation and resource for proteome-scale protein expression. Nat Genet 34, 35–41 (2003).

